# AsmMix: A pipeline for high quality diploid *de novo* assembly

**DOI:** 10.1101/2021.01.15.426893

**Authors:** Pei Wu, Chao Liu, Ou Wang, Xia Zhao, Fang Chen, Xiaofang Cheng, Hongmei Zhu

## Abstract

In this paper, we report a pipeline, AsmMix, which is capable of producing both contiguous and high-quality diploid genomes. The pipeline consists of two steps. In the first step, two sets of assemblies are generated: one is based on co-barcoded reads, which are highly accurate and haplotype-resolved but contain many gaps, the other assembly is based on single-molecule sequencing reads, which is contiguous but error-prone. In the second step, those two sets of assemblies are compared and integrated into a haplotype-resolved assembly with fewer errors. We test our pipeline using a dataset of human genome NA24385, perform variant calling from those assemblies and then compare against GIAB Benchmark. We show that AsmMix pipeline could produce highly contiguous, accurate, and haplotype-resolved assemblies. Especially the assembly mixing process could effectively reduce small-scale errors in the long read assembly.

## 1 Introduction

Genome assembly is the process of determining genome sequences, and a high-quality assembly is fundamental for understanding the biology of given species and important for downstream analysis. One can break down the quality of an assembly into several aspects. First, the assembly should be less fragmented, which can be evaluated by contig or scaffold N50. Second, there should be fewer error in the assembly, which can be further broken down into single-base accuracy, typically measured by Q-value (QV) and large-scale assembly errors, so-called misassemblies. Third, for a polyploid genome, the assembly should be haplotype-resolved, i.e., there should be one assembly representing each haploid.

A large number of laboratory and computational approaches have been developed to address the challenge of assembly. Next-generation sequencing (NGS) is a fast, low-cost, and high throughput technology. However, it is only capable of generating reads typically range in size 100~200bp, and with only shortrange information, which limits its ability to resolve repetitive regions. Singlemolecule sequencing (SMS) technologies, including Pacific Biosciences and Oxford Nanopore, produces reads with lengths varied from several kilobases to several millions of bases [1, 2, 3, 4], relatively high error rate (2~15%) [5, 6, 7]and high cost. While most errors can be corrected by taking consensus, still a large number of errors remain. The process of correcting those errors by NGS reads are always referred to as polishing, which typically starts from mapping NGS reads to SMS assemblies and correct errors by consensus or performing local assembly [8, 9, 10, 11].

Co-barcoded NGS is an augmented NGS technology, which starts from breaking chromosomes into millions of long DNA fragments and then tags reads from one fragment with the same barcode. Thus reads with the same barcode have a high probability to be derived from the same long DNA fragment, which means their positions on the chromosome should be within the length scale of DNA fragments and on the identical haplotype. Co-barcoding technology includes 10x Linked-read [12], stLFR [13] and a few others [14]. This technology could help us resolve repetitive regions and phase haplotypes. As a comparison, with only traditional pair-end NGS reads one would generate highly fragmented assemblies, typically with contig N50 ranges from 10~90Kb and scaffold N50 slightly larger than that [15, 16, 17]. But with co-barcoded reads, one could achieve haplotype-phased assemblies with contig N50 as long as 100~200Kb and scaffold N50 at 20~40Mb [18].

In this paper we demonstrate an assembly pipeline *AsmMix* which is optimized for the combination of high coverage co-barcoded, NGS and SMS reads. First, we build assemblies from co-barcoded and SMS reads independently. Then we perform the *assembly mixing* to correct the errors in the SMS assembly using co-barcoded assemblies. The strategy for the mixing is as follows: first, we compare two assemblies, consider their differences in small scale as errors in SMS assemblies, and then correct those errors by replacing with sequences in co-barcoded assemblies. The co-barcoded assemblies have a high scaffold N50, which is favorable for anchoring sequences to SMS assemblies.

## 2 Method

The AsmMix pipeline requires stLFR, NGS and SMS reads as input, the steps of AsmMix pipeline were shown in Fig.1 (a). In the first step, the AsmMix pipeline runs stLFRdenovo [19] on stLFR reads to generate a set of haplotype-resolved assemblies, and runs NECAT [20] on SMS reads and then polish by Pilon [8] using NGS reads to generate another assembly. The next step involves mixing these two into one. The goal of this step is to fix short-range errors in the SMS assembly using the stLFR assembly. First the AsmMix pipeline compares two assemblies to obtain a list of alignment blocks by running QUAST [21, 22] with SMS assembly as the target and stLFR assembly as the query. The QUAST software aligns two assemblies using minimap2 [23] and selects an optimal set of alignment blocks. Then the AsmMix pipeline further screens the set to remove possible inconsistencies (Fig.1 (b)). For each alignment block, the AsmMix pipeline performs base-level pairwise alignment by minimap2 and inspects the substitutions/indels between two sequences. If a substitution/indel has length smaller than a certain threshold (50 bases by default), it is considered as an error in the SMS assembly and replaced with sequence in the stLFR assemblies, substitutions/indels with length larger than the threshold are discarded. Using this strategy, most short-scale errors in the SMS assembly are corrected. This step is implemented in a python script. Each haplotype from stLFR assembly is mixed with SMS assembly independently to retain two haplotypes. In the following we describe inconsistent alignment blocks filtering and sequence replacement in detail.

**Figure 1:**
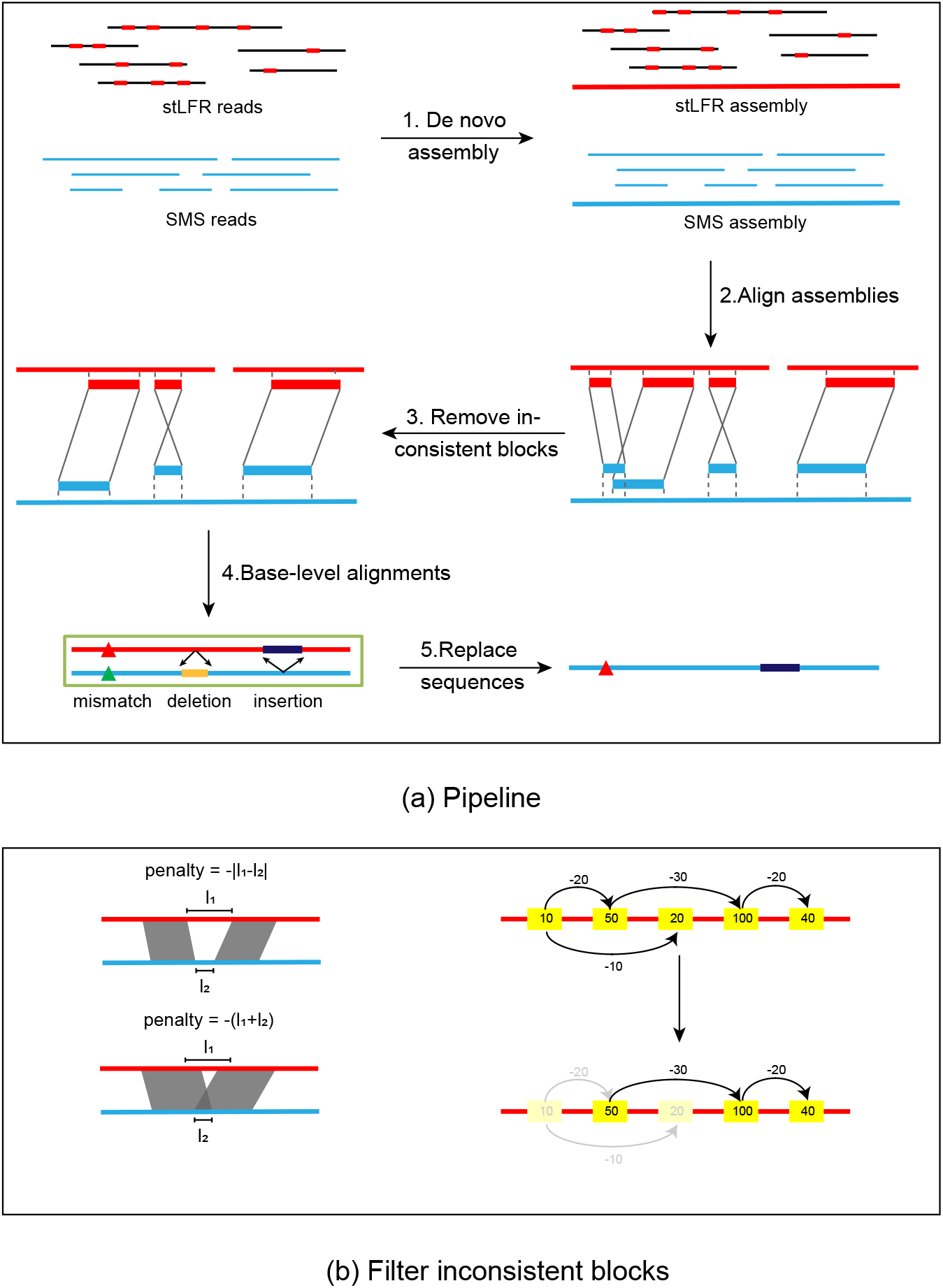
(a): An overview of pipeline AsmMix (b): Filter inconsistent blocks. In the figure left, we show an example of filtering inconsistent blocks, each yellow rectangle indicates an alignment block, the number in each rectangle indicates the length of the alignment block, and the number on the arrow indicates the penalty score. We seek a path that maximizes the sum of lengths and penalties it passes through. In the figure below, the shallow yellow rectangles indicate discarded alignment blocks, yellow rectangles, and black lines indicated selected blocks and paths.

### 2.1 Filter Alignment Blocks

While running QUAST the AsmMix pipeline set SMS assembly as the target and stLFR assemblies as the query. For each stLFR scaffold, a set of alignment blocks are generated by QUAST, sorted by aligned region on stLFR assembly and denoted by *b_i_, i* = 1,…, *n*. A penalty function *p(b_i_,b_j_*) for a pair of alignment blocks *b_i_, b_j_* is defined as follows: the penalty is set to —∞ if they map to distinct SMS contig, or their strands are different, or the length of their overlap on SMS scaffold exceeds 0.8 time of length of the shorter aligned region. Otherwise, the penalty is defined as follows: if their aligned regions on SMS scaffolds do not overlap, the penalty is defined as —|*l*_1_ — *l*_2_|, in which *l*_1_ and *l*_2_ are gap lengths between aligned regions on stLFR and SMS scaffolds respectively. And if aligned regions on SMS are overlapping, the penalty is defined as — (*l*_1_ + *l*_2_), in which *l*_1_ is gap length between aligned regions on stLFR scaffold and l_2_ is the length of overlap on SMS scaffold.

For a chain of alignment blocks {*b_k_*_1_, …, *b_k_*_m_}, *k*_1_ < … < *k*_m_. The score is defined as:

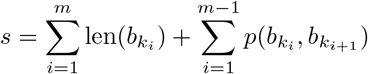

In which len(*b*) is the length of the aligned region on stLFR scaffolds. To select a chain that maximizes *s* we apply a dynamic programming algorithm: let s_*i*_ be the maximal score for chains terminated at b_*i*_, then s_*i*_ can be decided via this recursive relation:

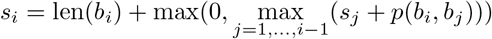

Thus if *s_i_* is maximal, the optimal chain must be terminated at *b_i_* and the whole chain can be recovered by a trace-back process. During implementation, we perform a two-round iteration for positive/negative strand.

After deciding an optimal chain, the alignment blocks in it are on the same target scaffold and the same strand. And for alignments blocks to different target scaffolds or different strands, we will include those back in the sequence replacement step.

### 2.2 Sequence Replacement

After filtering, the AsmMix pipeline clusters alignment blocks by SMS scaffold and sort alignment blocks on each SMS scaffold by their starting positions. For each SMS scaffold, the AsmMix pipeline looks through the alignment blocks to eliminate overlaps between every two consecutive regions based on the following rules: 1) if one region covers the next region, the latter will be deleted, 2) if two regions are overlapping, their overlaps will be assigned to the latter one. After dealing with overlapping parts, all selected regions on each SMS scaffold are independent and sent to minimap2 [23] with counterparts from their respective alignment blocks to do a pairwise alignment. Then the AsmMix pipeline looks into alignment details by parsing cs tags in paf [24] files. The final replacement will ignore N bases in stLFR assemblies, and there is a length threshold (50 bases by default) to control whether to replace when there is an indel signal in the cs tag. Then the AsmMix pipeline concatenates the replaced sequences and sequences from regions not covered by alignment blocks as the final output.

## 3 Results

We tested the AsmMix pipeline on a dataset consisting of stLFR reads (~84x), MGI PCR-free reads (~120x), and Oxford Nanopore Technology (ONT) reads (~120x) of NA24385. The longest 50x ONT reads were extracted to perform NECAT assembly. All the assemblies were evaluated by QUAST and primate gene marker set *primates_odb10* by BUSCO [25, 26]. The results were enlisted in Table 1. Most metrics do not change significantly after mixing, except the number of indel decreases from 0.39 to 0.31 per 1Kbp, which reflects a decrease of indel errors by the mixing process.

**Table 1:** Quality assessment of *de novo* assemblies of NA24385. stLFR hap1/2 is the haplotype-resolved assembly built from stLFR_denovo, Necat is the assembly build by Necat and polished by Pilon, Mix_hap1/2 is the haplotype-resolved assembly after assembly mixing. Total length, scaffold NG50, contig NG50, genome fraction, duplication ration, scaffold NGA50, contig NGA50, number of misassemblies, mismatch per 1Kbp, and indel per 1Kbp were evaluated by Quast v5.0.2 with reference genome *hs37d5*, and parameters “*-s -m 10000 - x7000*”. The value of row Total length, Genome fraction, and Duplication ratio before the slash are values of scaffolds, and the values after the slash are values of contigs. BUSCO complete ratio was evaluated by BUSCO v4 with marker set *primates_odb10*.

|  | stLFR_hap1 | stLFR_hap2 | Necat | Mix_hap1 | Mix_hap2 |
| --- | --- | --- | --- | --- | --- |
| Total length | 2,899,262,788/<br>2,624,851,442 | 2,899,974,893/<br>2,623,423,014 | 2,880,654,213 | 2,880,459,545 | 2,880,632,458 |
| Scaffold NG50 | 21,283,539 | 21,282,592 | - | - | - |
| Contig NG50 | 85,008 | 84,928 | 34,759,314 | 34,757,611 | 34,759,622 |
| Genome fraction | 90.688%/<br>90.865% | 90.636%/<br>90.806% | 97.815% | 97.814% | 97.810% |
| Duplication ratio | 1.058/<br>1.004 | 1.059/<br>1.004 | 1.009 | 1.008 | 1.009 |
| Scaffold NGA50 | 1,826,678 | 1,826,238 | - | - | - |
| Contig NGA50 | 82,847 | 83,028 | 15,981,976 | 15,594,006 | 15,594,501 |
| #Misassemblies | 4,519 | 4,493 | 2,536 | 2,548 | 2,539 |
| Mismatch per 1Kbp | 1.0174 | 1.0128 | 1.2765 | 1.2888 | 1.2867 |
| Indel per 1Kbp | 0.2504 | 0.2494 | 0.3932 | 0.3116 | 0.3114 |
| BUSCO complete ratio | 86.3% | 86.1% | 89.4% | 89.9% | 89.9% |

To further analyze the accuracy of assembly, we performed variant-calling and compare results against a well-established benchmark: the Genome in a Bottle benchmark (small variant v3.3.2, structural variant v0.6). The accuracy of variant calling reflects the accuracy of assembly. The variant-calling from assemblies was performed by minimap2 and paftools. For haplotype-phased assemblies, variations were called separately and combined using vcftools [27]. Evaluations against benchmarks were performed by rtgtools [28] for small variants and truvari [29] for structural variants. We evaluated phasing accuracy by the short/long switch error rate and phase block N50. The short switch error is the error that flips a single heterogeneous site, and the long switch error flips the haplotype of all heterogeneous sites after a certain position. To count short/long switch errors, we minimize the penalty function *5_n_long__* + *n_short_*, in which *n_short_* is the number of short switch errors and *n_long_* is the number of long switch errors. Short/long switch error rate is the ratio of the number of short/long switch errors to the number of common heterogeneous SNPs between the benchmark and call set. Phase blocks are defined as regions cutting by long switch errors, and phase block N50 is the N50 of their lengths.

Those results are enlisted in Table 2. For the SNP, mixing leads to a 3.8/3.8- fold decrease of false negatives, and a 3.4/1.4-fold decrease of false positives, with/without ignoring haplotyping difference. For the indel, mixing leads to a 3.0/3.4-fold decrease of false negatives, and a 3.6/4.8-fold decrease of false positives, with/without ignoring the haplotyping difference. For the large SVs, the false negative for insertion/deletion remains almost unchanged, but the false positives display a 1.8/1.6-fold increase. Our test shows that the mixing procedure could significantly improves the accuracy of short variant, which means a large proportion of small-scale errors are corrected. We observed a decrease in the performance of long variant calling, which could partially be explained by the fact that short reads could not resolve repetitive regions effectively. And all metrics for phasing display a slight improvement when comparing with stLFR assemblies.

**Table 2:** Evaluation of assembly by variant calling. In the row SNV, short variations of call and benchmark are separated into SNP and indel by vcftools and compared by rtgtools separately. The value before the slash are without and after the slash are with the parameter “–squash-ploidy”, which allow matches ignoring the haplotyping difference. In the row SV, call and benchmark are compared by truvari with the parameter “-r 1000 –passonly” and false-positive insertion/deletion were counted by a perl script, in which variants with reference sequence longer than alternative sequence are counted as deletions and otherwise insertions, variants with “N” in the alternative sequence are ignored. In the phasing column, common heterogeneous SNP was selected and their phases were compared, phased SNP ratio is the ratio of common heterogeneous SNP against heterogeneous SNP in the benchmark, short switch error is defined as error of flipping phase of a single variant, and long switch error defined as error of flipping all phase after of a variant. The penalty for long switch error is 5 times of penalty for switch error and minimized by a dynamic programming scheme. Phase blocks are defined as regions cut by the location of long switch error.

| variation type | variation subtype | Metric | stLFR | NECAT | Mix |
| --- | --- | --- | --- | --- | --- |
| SNV | SNP | TP | 2,330,486/2,523,914 | 1,161,996/2,067,723 | 2,536,637/2,779,428 |
|  |  | FN | 698,572/505,144 | 1,867,062/961,337 | 492,421/249,635 |
|  |  | FP | 253,797/60,369 | 975,393/69,669 | 291,137/48,351 |
|  |  | Precision | 0.9018/0.9766 | 0.5437/0.9674 | 0.8970/0.9829 |
|  |  | Sensitivity | 0.7694/0.8332 | 0.4498/0.6826 | 0.8374/0.9176 |
|  | Indel | TP | 334,312/377,364 | 162,087/284,873 | 358,833/408,144 |
|  |  | FN | 125,053/82,001 | 297,278/174,492 | 100,541/51,200 |
|  |  | FP | 72,872/29,827 | 355,587/232,840 | 97,589/48,279 |
|  |  | Precision | 0.8210/0.9268 | 0.3131/0.5502 | 0.7862/0.9842 |
|  |  | Sensitivity | 0.7278/0.8215 | 0.3529/0.6201 | 0.7811/0.8885 |
| SV | Insertion | TP | 1811 | 3752 | 3906 |
|  |  | FN | 3631 | 1779 | 1659 |
|  |  | FP | 539 | 343 | 627 |
|  |  | Precision | 0.7707 | 0.9162 | 0.8616 |
|  |  | Sensitivity | 0.3329 | 0.6784 | 0.7018 |
|  | Deletion | TP | 2750 | 2420 | 2539 |
|  |  | FN | 1449 | 1690 | 1537 |
|  |  | FP | 5983 | 763 | 1223 |
|  |  | Precision | 0.3148 | 0.7603 | 0.6749 |
|  |  | Sensitivity | 0.6548 | 0.5888 | 0.6229 |
| Phasing |  | Phased SNP ratio | 0.6881 | - | 0.7224 |
|  |  | Short switch rate | 0.0515 | - | 0.0502 |
|  |  | Long switch rate | 0.0008 | - | 0.0007 |
|  |  | Phase block N50 | 4,802,248 | - | 5,015,779 |

### 3.1 Computational cost

The AsmMix pipeline were tested on the dataset described above with a 32-CPU computational node with 3TB memory. In the pipeline, Necat takes 7 days, Pilon polishing takes 2 days and stLFR_denovo takes 4 days, the mixing step can be finished within 2 hours, with most of the time spent by QUAST. The whole pipeline takes around 14 days to complete.

## 4 Discussion

It is common practice to perform assembly with a combination of data from different technologies, but the problem of what is the optimal strategy for data combination is still wide open. In this paper we propose an assembly pipeline that is capable of reaching three goals: contiguity, single-base accuracy, and haplotype-resolved, making full use of high coverage stLFR, NGS and SMS reads. The accuracy of assembly is enough for assembly-based SNP, InDel and structure variant calling with competitive performance.

Moreover, as assembly mixing is implemented as an independent module, the AsmMix pipeline is compatible with any assemblers for SMS and co-barcoded reads, this could allow users to test and customize their pipeline in a more flexible manner.

Our plan for the AsmMix pipeline includes the following directions. First we use only co-barcoded reads to generate longer contigs in co-barcoded reads assemblies and correct short-range errors, but it is expected that with that information one could further scaffold SMS assemblies. Most of the scaffolding methods start from mapping reads to SMS assemblies, which takes large computational resources [30, 31], we will design the method of scaffolding using co-barcoded reads assemblies. Second, we only use co-barcoded reads in phasing, in the future we will study how to integrate SMS reads to improve phasing performance.

## 5 Availability of data and materials

The AsmMix pipeline is available on GitHub (https://github.com/BGI-biotools/AsmMix). The stLFR co-barcoded reads of HG002 is available at https://db.cngb.org/search/run/CNR0026818/. We got 120X Nanopore long reads of HG002 from Oxford Nanopore Technologies.

## Notes

### Competing Interest Statement

The authors have declared no competing interest.

